# Single-Particle STORM Imaging Quantifies Coating Heterogeneity Among Cell Membrane-Coated Nanoparticles

**DOI:** 10.64898/2026.09.28.755175

**Authors:** Jimmy Blauser-Wilson, Ali Zareein, Sao Puth, Shruti Sunil Jadhav, Raymond Chen, Abigail M Hernandez, Andrea Joseph, Jialiu Zeng, James L Hougland, Yaoying Wu

**Affiliations:** Department of Biomedical and Chemical Engineering, Syracuse University, Syracuse, NY 13210, USA; The BioInspired Institute for Material and Living Systems, Syracuse University, Syracuse, NY 13210, USA; The Global Health Research Center, University of Health Sciences, Khan Daun Penh, Phnom Penh, 12000, Cambodia; College of Science, University of Texas at San Antonio, San Antonio, TX, 78249, USA; Department of Chemistry, Syracuse University, Syracuse, NY, 13244, USA; Department of Biology, Syracuse University, Syracuse, NY, 13244, USA; Department of Microbiology & Immunology, SUNY Upstate Medical University, Syracuse, NY 13210, USA

**Keywords:** cell membrane-coated nanoparticles, STORM imaging, antigen presentation, dendritic cell membrane-coated nanoparticles, membrane coating coverage, membrane coating composition

## Abstract

Cell membrane-coated nanoparticles (CNPs) are a potentially transformative biomimetic targeted-delivery platform, which can engage therapeutic targets through source-cell membrane protein functions or homotypic interactions. However, their therapeutic development is limited by the current inability to accurately resolve membrane coating completeness, and CNP particle-to-particle variation. Here, we introduce a STochastic Optical Reconstruction Microscopy (STORM)-based approach that resolves population composition and quantifies membrane coating coverage at the single-particle level. Applying it to dendritic cell membrane- and HeLa cell membrane-coated nanoparticles (DCmPs and HeLamPs), we revealed substantial heterogeneity within both particle types and a cell-type dependence of coating outcomes, corroborated by confocal imaging and flow cytometry. We also quantified membrane protein composition, including peptide major histocompatibility complex class I (pMHC-I) among DCmPs, and confirmed that DCmPs preferentially engaged antigen-specific T cells in coculture. This single-particle framework establishes a quantitative standard for CNP characterization, providing a foundation for quality control and rational design.

**TABLE OF CONTENT:** 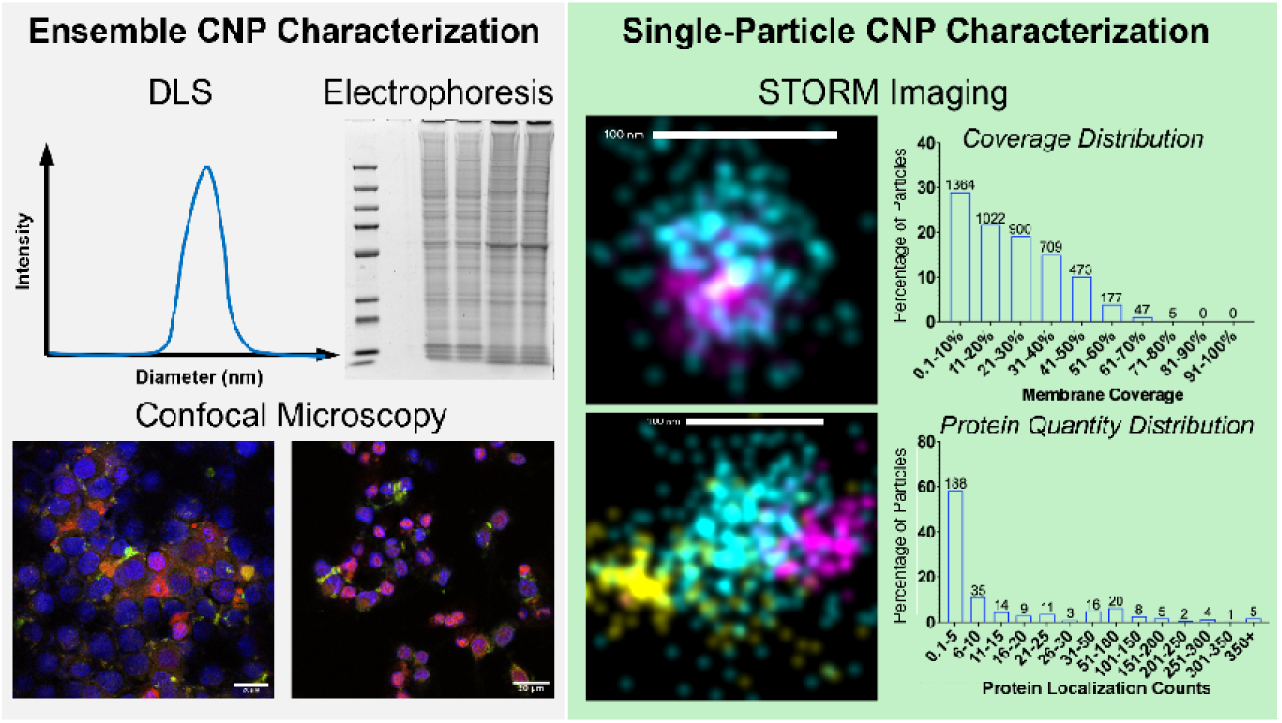

A single-particle super-resolution imaging method is developed to determine the coating composition and completeness across membrane-coated nanoparticles. Examining two types of coated nanoparticles reveals striking variations in coating coverage and protein composition, and shows cell type-dependent coating quality. This technology provides a new detailed and precise quantality control for membrane-coated nanoparticle design.

## MANUSCRIPT

Cell membrane-coated nanoparticles (CNPs) integrate synthetic particulate cores with cell-derived membrane coating, representing a potentially transformative biomimetic drug delivery strategy.[1–3] The membrane coating endows CNPs with biological function resembling the membrane from source cells, thereby enabling a broad range of applications. For instance, cancer cell membrane-coated nanoparticles have been extensively studied for their cancer-targeted drug delivery ability owing to the high affinity homotypic binding between CNPs and the source cells.[3–6] Red blood cell (RBC) membrane-coated nanoparticles inherit immune evasion capability from RBCs, resulting in prolonged circulation and improved biodistribution for cancer targeting.[7–9] Membranes of immune cells have also been leveraged for the CNP system.

Cytokine receptors and adhesive molecules from macrophage or neutrophil membranes enable CNPs to extract inflammatory cytokines for anti-inflammation treatment, or to detect circulating cancer cells.[10–15] More recently, dendritic cell membrane-coated nanoparticles (DCmPs) have been developed to facilitate antigen-specific T-cell-targeted therapy, owing to the cognate interaction between major histocompatibility complex (MHC) and T-cell receptor (TCR).[16–18] Various types of particles have been explored as CNP particulate cores, including poly(lactic-co-glycolic acid) (PLGA) nanoparticles (NPs), mesoporous silica nanoparticles (MSN), and gold NPs, facilitating the encapsulation of a wide range of pharmaceutical or imaging agents.[19,20]

CNPs are conventionally produced through sonication, extrusion, or a combined (sonication followed by extrusion) approach to mechanically induce association between isolated membrane proteins and particulate cores.[3,18,21] Although the combined coating approach improves CNP uniformity at a population level relative to sonication and extrusion, all three coating approaches yield highly heterogeneous final products, consisting of CNPs, noncoated particulate cores, and membrane proteins.[18] Beyond population composition heterogeneity, the membrane coverage (the percentage of particle surface covered by membrane proteins) likely varies between coating approaches and likely substantially across individual CNPs.[21,22] The CNP heterogeneity inevitably introduces variation within and between batches, which potentially undermines the translational potential. Detailed characterization of the coating products at both population and single-particle levels is therefore crucial in guiding the rational development of CNP-based therapeutics.

Ensemble characterization approaches, such as Dynamic light scattering (DLS) and gel electrophoresis assays, such as western blot, have been extensively utilized to confirm population level membrane protein association and protein quantity estimation, but largely fail to resolve particle-to-particle variation. To address this limitation, researchers have developed indirect methods to assess coating completeness. Zhang et al. synthesized biotinylated PLGA cores for membrane coating when they observed aggregation of CNPs in streptavidin protein-containing buffer. Their finding indicates biotin-PLGA cores remain accessible, suggesting CNP coating is incomplete.[23] Vesa-Pekka and coworkers developed a CNP system with MSN cores that contain nitro-2,1,3-benzoxadiazol-4-yl (NBD) fluorescent molecules. Since the fluorescent NBD can be quenched through a reduction reaction by dithionite (DT), the extent of NBD quenching can be correlated with CNP membrane coverage. Complete membrane coating will prevent DT from quenching NBD fluorescence, but CNPs that are partially coated or noncoated will be quenched by DT treatment.[21,22] We and others leveraged homotypic interactions between CNPs and source cells and developed cellular uptake assays to determine CNP composition.[4,18,24] During the interactions, the source cells are treated with CNPs formed by using distinctly fluorescently labeled particulate core and membrane protein. The CNP composition can thus be estimated based on the fluorescence(s) of particles internalized by source cells using confocal microscopy or flow cytometry. Flow cytometry has additionally been employed to directly assess the CNP membrane protein composition, although the accuracy can be limited by the difficulty in establishing single-particle resolution for nanoparticle samples.[18] However, all these characterization approaches are largely unable to distinguish individual particles or quantify protein composition at the single-particle level. This critical knowledge gap hinders the development of CNP technologies, especially the platforms that rely on membrane protein-dependent functions.

To address this knowledge gap, we developed a STochastic Optical Reconstruction Microscopy (STORM) imaging-based strategy for single-particle characterization of CNP membrane coating.[25–29] We applied this approach to determine the particle composition and membrane coverage of two types of CNPs: DCmP (PLGA NPs coated with membrane from DC2.4 cells, murine DC cell line) and HeLamP (PLGA NPs coated with membrane from HeLa cells, human cervical cancer cell line) at a single-particle level, benchmarked against two existing orthogonal approaches, cell uptake assay and flow cytometry. Additionally, because DCmPs carry key proteins to engage antigen-specific T cells, including MHC class I (MHC-I), CD86, and ICAM-1, we quantified the composition of these proteins, particularly the SIINFEKL/MHC-I complexes, using STORM imaging.[18] We further confirmed antigen-specific T cell binding in a T cell co-culture model.

Cell membrane proteins were isolated from HeLa cells or OVA/LPS-stimulated DC2.4 cells using Dounce homogenizer and were confirmed to preserve the characteristic protein features of source cells via SDS-PAGE gel. (Fig S1) We subsequently produced DCmPs and HeLamPs using the corresponding membrane proteins via a combined (sonication followed by extrusion) coating approach.[18] (Fig 1A) Both DCmPs and HeLamPs show increased diameters (DCmP: 140.40nm; HeLamP: 144.13 nm) relative to bare PLGA NPs (120.75 nm) according to ZetaView. (Fig 1B) Polydispersity index (PDI) of DCmP was measured at 0.31, HeLamP at 0.44, and bare NPs at 0.52. (Fig 1C) Surface charge of DCmPs and HeLamPs also confirms they are associated with proteins as the zeta potential of DCmPs and HeLamPs decreases to -33.20 mV and -28.69 mV, respectively, from -18.79 mV (PLGA NPs). (Fig 1D) DLS measurement also confirmed the size increase and surface charge decrease for both DCmPs and HeLamPs. (Fig S2A, S2B, S2C) Collectively, these data confirm the presence of protein coating on PLGA NPs.

**Figure 1.**
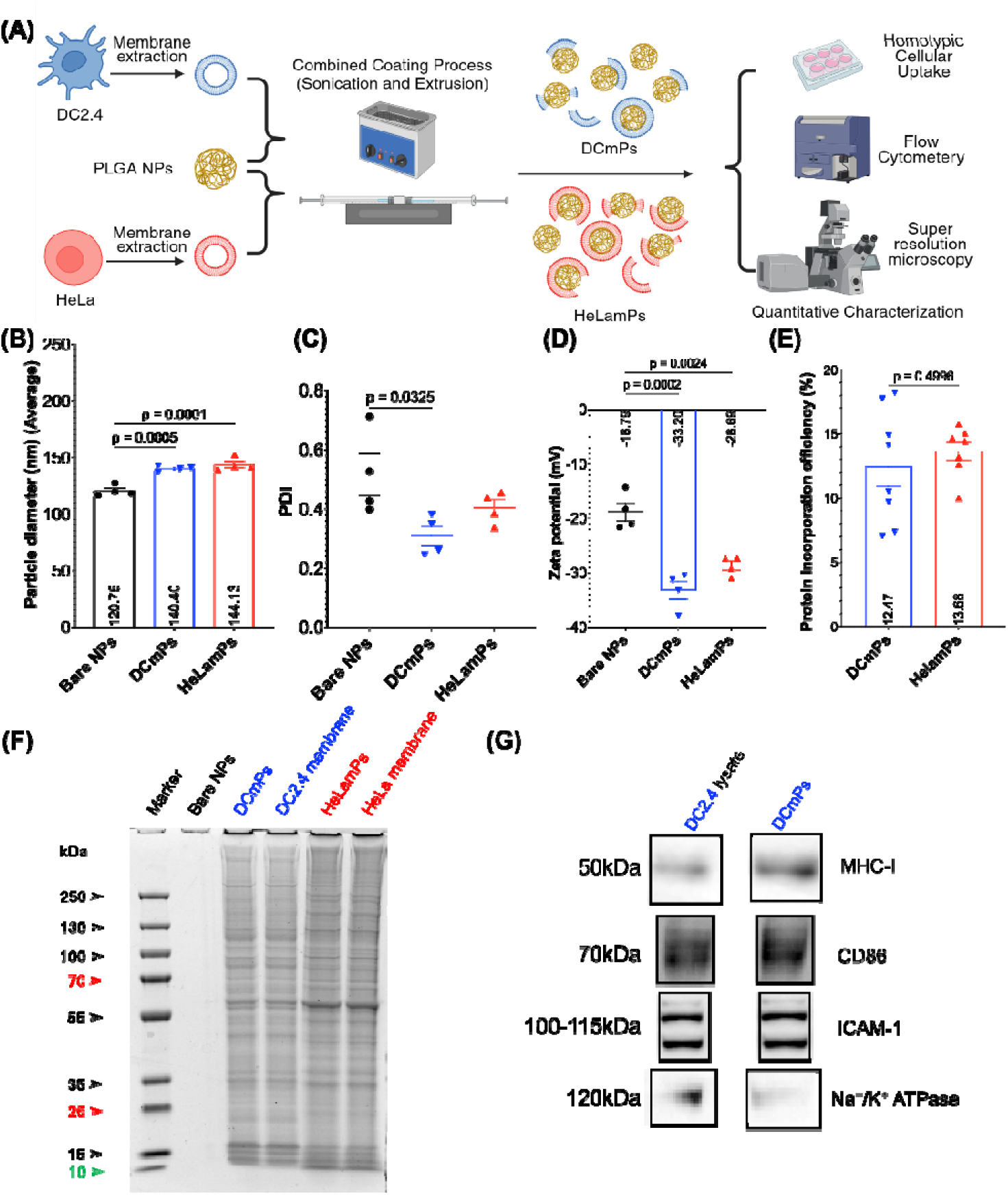
DCmPs and HeLamPs carry protein coatings resembling source cell membrane proteins. (A) Overview schematics. Cell membranes are isolated from DC2.4 or HeLa cells and coated onto PLGA NPs using a combined coating process (sonication followed by extrusion). The obtained DC2.4 cell membrane-coated nanoparticles (DCmPs) and HeLa cell membrane-coated nanoparticles (HeLamPs) are characterized using several approaches, including a homotypic cellular uptake assay, flow cytometry, and super-resolution microscopy. The particle diameters (B) polydispersity index (PDI) (C) and zeta potentials (D) of DCmPs and HeLamPs were determined using ZetaView particle analyzer. Statistical significance is determined using one-way ANOVA following Tukey’s multiple comparison test. N=4. (E) The protein incorporation efficiencies of the coating process for DCmPs and HeLamPs were determined using Bradford assay. P value is determined using unpaired t test with Welch’s correlation. N=8. (F) Protein profiles of membrane-coated nanoparticles and the cell membrane proteins isolated from each type of source cells were determined using SDS-PAGE gel analysis. (G) The presence of antigen presentation-related proteins in both DC 2.4 cell lysate and DCmPs was confirmed using western blot gel analysis.

To determine the protein incorporation efficiency (the fraction of total proteins attached to CNPs), we quantified the amount of protein in PBS before and after the coating process via Bradford Assay. 12.5% of membrane proteins were coated onto DCmPs, while HeLamPs have a slightly higher protein incorporation efficiency at about 13.7%. (Fig 1E) SDS-PAGE gel also confirmed that the protein profiles of DCmPs and HeLamPs resemble the respective membrane proteins. (Fig 1F) Additionally, we confirmed that DCmPs carry several key proteins for cognate DC-T cell interaction, including MHC-I, CD86, ICAM-1, and Na+/K+ ATPase, using western blot. (Fig 1G) Furthermore, both DCmPs and HeLamPs were colloidally stable in PBS for 24 hours at 4 °C, as their diameters remained stable during the experimental period. (Fig S3A, S3B, S3C, S3D, S3E, S3F)

We and others have utilized homotypic interaction-mediated cellular uptake to determine the composition of the final coating product.[4,18,24] Here, we aim to confirm this homotypic interaction-based approach and determine the CNP coating efficiency, which is defined as the percentage of PLGA NPs that are coated with membrane proteins. To this end, we produced both DCmPs and HeLamPs using Rhodamine B (RhoB)-encapsulated PLGA NPs and CFSE-stained membrane proteins of DC2.4 cells and HeLa cells respectively. After confirming that the diameters and PDI of both fluorescently labeled DCmPs and HeLamPs are consistent with their unlabeled counterparts. (Fig 1B, 1C, S4) We treated DC2.4 and HeLa cells with either particle.

The frequency of cells that took up CNPs (CFSE+RhoB+) was determined via flow cytometry after 2 and 4 h treatment. DCmPs were rapidly taken up by DC2.4 cells, with the uptake rate (CFSE+RhoB+) reaching 26.4% at 2 h and increasing to 51.2% at 4 h, while significantly fewer HeLamPs were taken up by DC2.4 cells (15.9% at 2 h and 36.3% at 4 h). (Fig 2A, S5) Similarly, HeLa cells also exhibited preference in uptake of HeLamPs over DCmPs. Two-hour incubation leads to 30.3% of uptake for HeLamPs, but only 1.9% for DCmPs. After 4 h incubation, 82.9% of HeLa cells were positive for HeLamPs, relative to 25.6% for DCmPs. (Fig 2B, S6) These results confirmed the rapid uptake of CNPs by the source cells and supports homotypic cell uptake approach for CNP composition analysis. The CNP coating efficiency can thus be estimated based on the uptake rate by the source cells, as 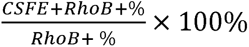, yielding 52.2% and 97.4% for DCmPs and HeLamPs respectively. (Table 1)

**Figure 2:**
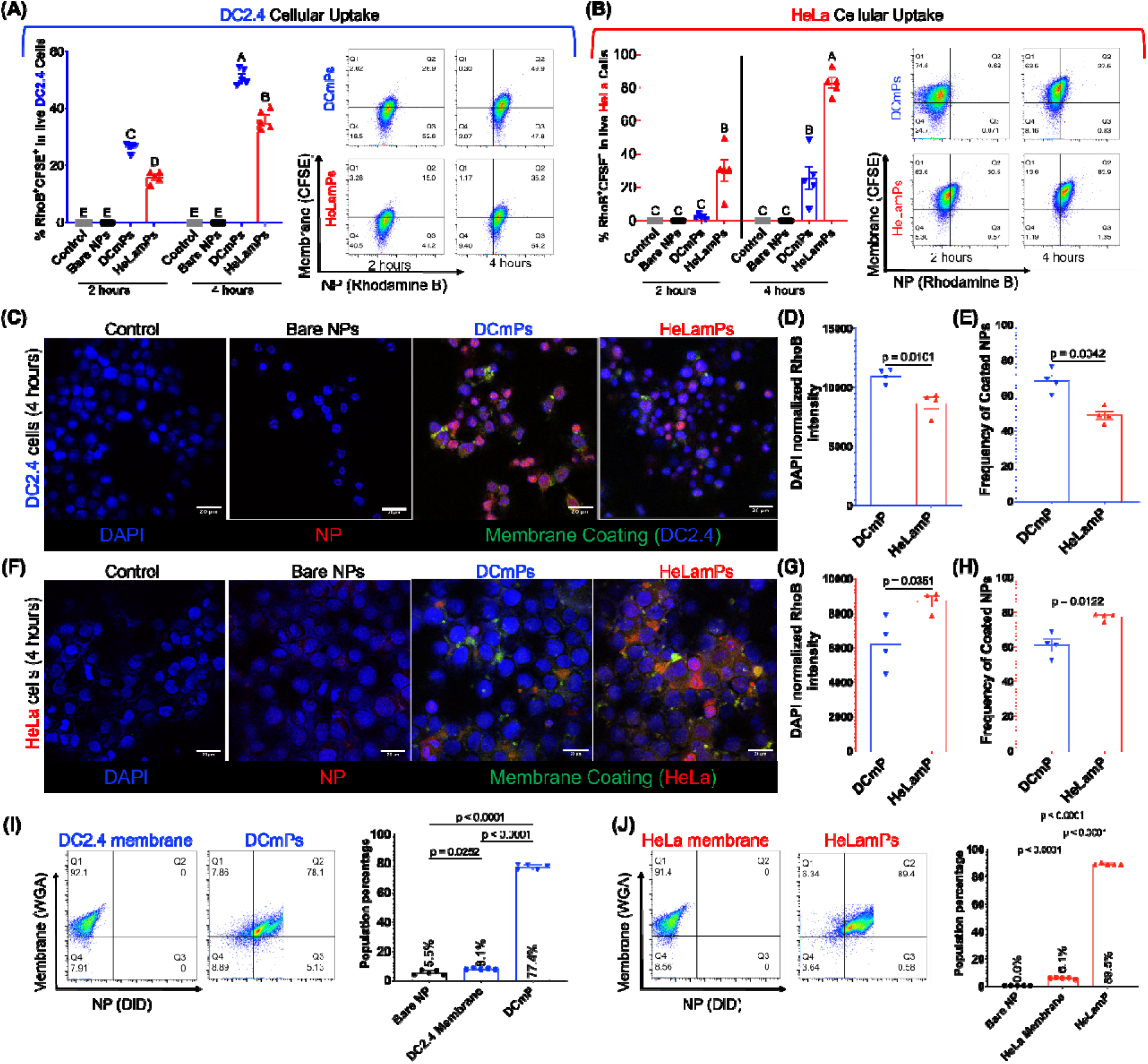
Coating efficiency Assessments for DCmPs and HeLamPs using homotypic cellular uptake assay and flow cytometry. DC2.4 cells (A) or HeLa cells (B) were incubated with fluorescently labelled DCmPs or HeLamPs for 2 or 4 hours. Particle cellular uptake was determined using flow cytometry (LEFT: Data summary; RIGHT: Representative flow plots). Statistical significance is determined using two-way ANOVA following Tukey’s multiple comparison test. Columns that do not share letters are significantly different (p < 0.05). N=5. DC2.4 cells (C) and HeLa cells (F) were cocultured for 4 hours with bare particles, DCmPs or HeLamPs. CNPs were labelled with CFSE (membrane coating) and RhoB (PLGA core). Nuclei were stained with DAPI. The particle internalizations were visualized using confocal microscopy. RhoB intensity normalized against DAPI area was determined to assess particle uptake by DC2.4 cells (D) or HeLa cells (G). The coating efficiencies were quantified based on the colocalization rate between CFSE and RhoB for DC2.4 cells (E) and HeLa cells (H). Statistical significance was determined using unpaired t-test, N=4. Coating efficiencies of DCmPs (I) and HeLamPs (J) were directly measured using flow cytometry, with CNP membranes stained by Alexa Fluor 488-conjugated Wheat Germ Agglutinin (WGA) and NP core stained with DiD. (LEFT: Representative images; RIGHT: Data summary) Statistical significance was determined using one-way ANOVA following Tukey’s multiple comparison test. N=4.

**Table 1.**
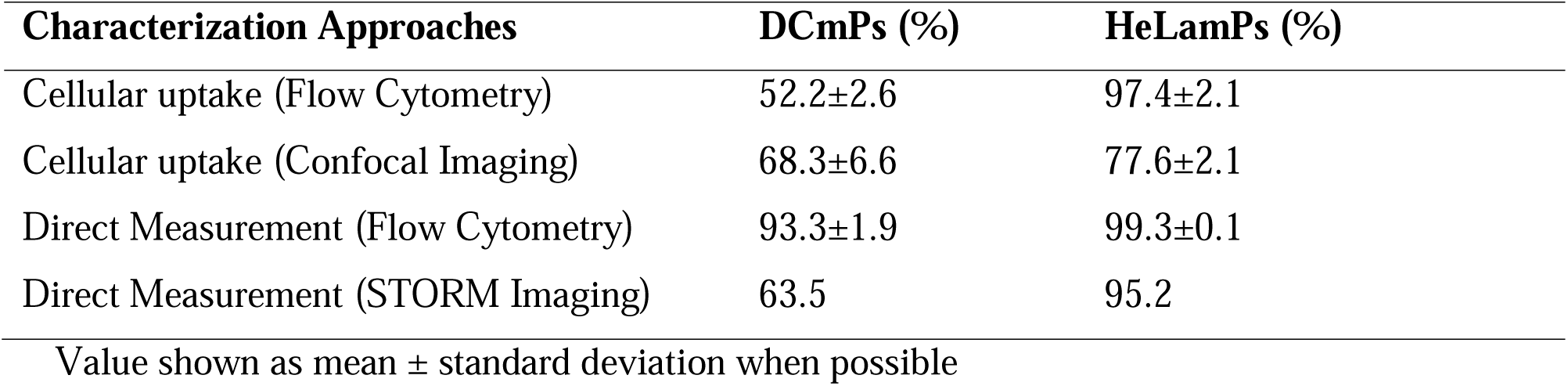
Summary of coating efficiency determined using different approaches.

| Characterization Approaches | DCmPs (%) | HeLamPs (%) |
| --- | --- | --- |
| Cellular uptake (Flow Cytometry) | 52.2±2.6 | 97.4±2.1 |
| Cellular uptake (Confocal Imaging) | 68.3±6.6 | 77.6±2.1 |
| Direct Measurement (Flow Cytometry) | 93.3±1.9 | 99.3±0.1 |
| Direct Measurement (STORM Imaging) | 63.5 | 95.2 |

Value shown as mean ± standard deviation when possible

We further leveraged this homotypic interaction to assess the coating frequency of CNPs using confocal imaging after 4 h of incubation. (Fig 2C, 2F, S7 and S8) Consistent with flow cytometry experiments, both source cells showed significantly higher uptake of their corresponding CNPs than of mismatched CNPs, indicated by the high DAPI-normalized RhoB (PLGA NPs) intensity in cells treated with their corresponding CNPs relative to mismatched CNPs. (Fig 2D and 2G) We determined the Manders’ colocalization coefficient for the overlap of CFSE (membrane) with RhoB for all treatment conditions. (Fig 2E and 2H) Taking the coefficient values from cells treated with corresponding CNPs as the CNP coating efficiencies, the coating efficiency was determined to be about 68.3% for DCmPs and about 77.6% for HeLamPs. (Table 1) HeLamPs show consistently higher coating efficiency than DCmPs between confocal imaging and flow cytometry measurements.

Although this homotypic cellular uptake assay circumvents the resolution limitation of confocal imaging by visualizing nanoparticles within cells, the accuracy in coating assessment using this approach relies on several critical assumptions: (1) that both CNPs and NP cores are taken up by the source cells equally; (2) that cellular uptake does not interrupt CNP structures. However, bare NPs alone are not internalized as efficiently as CNPs by cells. (Fig 2A, 2B, 2C, and 2F) Lysosomal activities also can disrupt membrane coating upon internalization. Thus, the accuracy of confocal imaging assessment is potentially suboptimal. We also carried out flow cytometry experiments to directly assess CNP coating frequency as an alternative strategy. Both DCmPs and HeLamPs were produced using DiD-encapsulated PLGA NPs and Alexa Fluor 488-conjugated Wheat Germ Agglutinin (WGA)-labeled membrane proteins, and the frequency of DiD+WGA+ population among DiD+ population is defined as coating frequency. For DCmPs, about 77.4% of the total particle populations are WGA+DiD+, resulting in a coating efficiency of 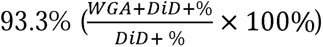. The frequency of HeLamPs among the total population is slightly higher at 89.5%, yielding a coating efficiency of 99.3%. (Fig 2E and 2H, Table 1). The results from direct flow cytometry measurement closely align with the value determined via homotypic cellular uptake assays, which HeLamPs showing higher coating frequency. However, because Fortessa flow cytometer is not designed for nanoparticle analysis, the potential suboptimal single-particle flow may undermine the accuracy of coating assessments.

Because of the accuracy limitation of confocal imaging and flow cytometry in determining CNP coating efficiency and their inability to resolve heterogeneity among individual particles, we adopted a STORM super-resolution imaging approach to visualize membrane coating and to quantify the coating frequency of DCmPs and HeLamPs.[30–32] To facilitate STORM imaging, we prepared these CNPs using RhoB-PLGA NPs and WGA-stained membrane proteins. The nanoparticles were captured onto poly-L-lysine-treated sample slides for STORM imaging, and were determined algorithmically based on the RhoB/WGA clustering within the particle diameters determined by ZetaView. STORM images revealed the colocalization between membrane proteins (teal) and PLGA NPs (magenta) at single-particle level, visually confirming the association between PLGA NPs and membrane proteins for both DCmPs and HeLamPs. (Fig3A and 3B) The presence of both empty protein vesicles and uncoated PLGA NPs in both DCmP and HeLamP samples was also confirmed. (Fig3A and 3B) Specifically, among all 8,149 particles captured in DCmP samples, 57.6% were coated nanoparticles (WGA+RhoB+, DCmPs) and 33.1% were bare PLGA NPs (WGA-RhoB+), yielding a coating frequency of 63.5% 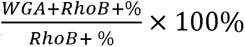, Fig 3C, Table 1). The remaining 9.1% consist of empty membranes. (Fig 3C) For HeLamP samples, we captured 4,049 particles via STORM imaging. Among them, 74.8% were HeLamPs, 3.8% were uncoated PLGA NPs, and 21.4% were lone membrane proteins, with a coating frequency of 95.2%. (Fig 3E and Table 1) Using this single-particle STORM imaging approach, we directly assessed CNP composition and confirmed that CNP coating efficiency is cell type-dependent and that HeLamPs consistently exhibit higher coating frequency than DCmPs.

**Figure 3:**
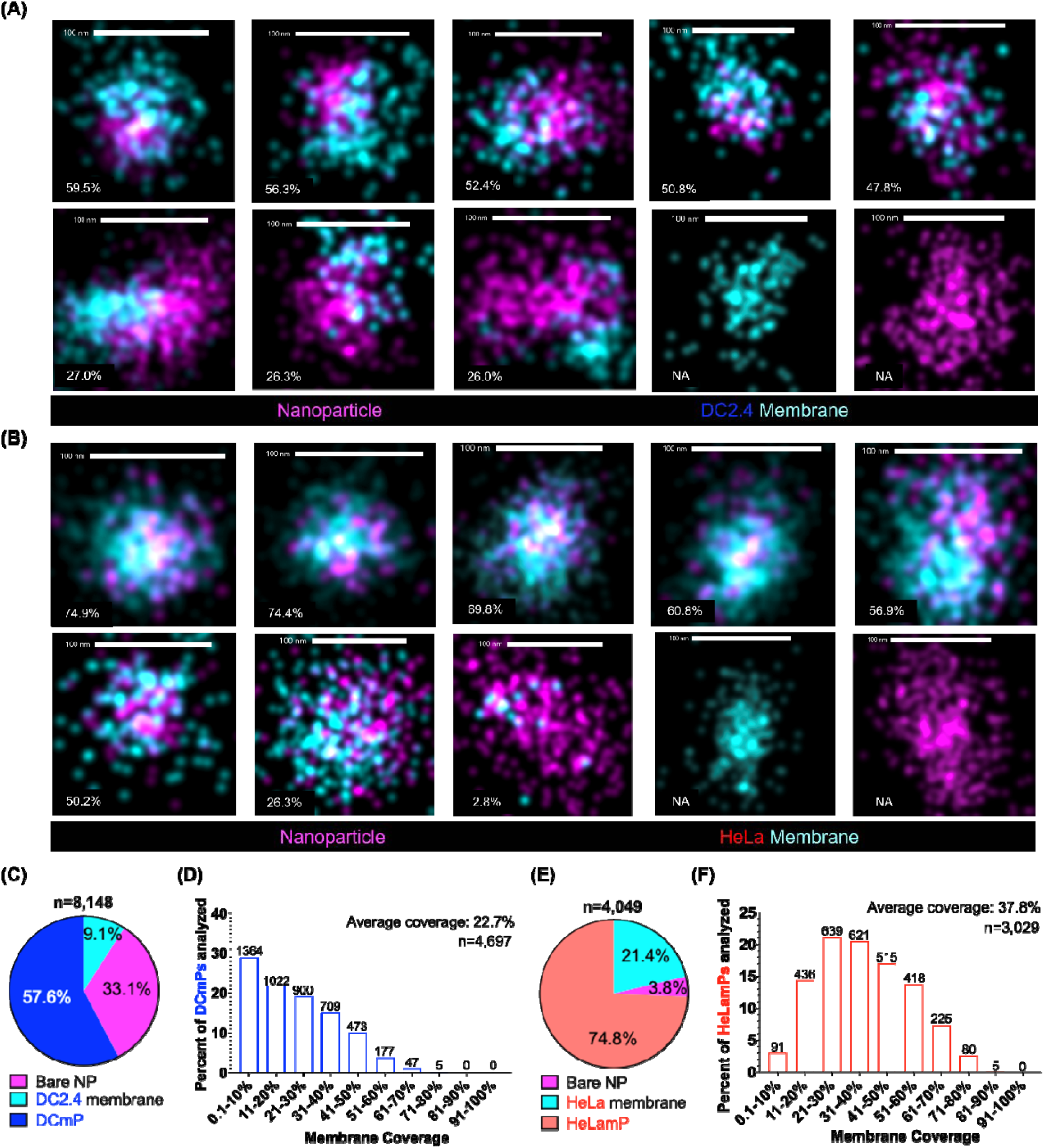
Single-particle analysis of membrane coverage of DCmPs and HeLamPs using super-resolution microscopy. DCmPs and HeLamPs were produced using RhoB-encapsulated PLGA NP cores, with lipid membrane stained with Alexa Fluor 488-conjugated WGA before imaging. Fluorescently stained particles were captured onto a Poly-L-Lysine-treated imaging chamber for STORM imaging. Representative images of single DCmPs (A) and HeLamPs (B) are presented. Percentage represents the membrane coverage of the corresponding particle. (C) The composition of all imaged DCmPs was summarized in pie chart. Of 8,149 particles imaged, 57.6% of particles were DCmPs, 33.1% were bare NPs, and 9.1% were empty membrane. (D) The frequency of the membrane coverage of all imaged DCmPs is summarized. The number above each bar represents particle numbers for each bin. (E) The compositions of all imaged HeLamPs are summarized in pie chart. Among the 4,049 imaged particles, 74.8% are HeLamPs, 3.8% bare NPs, and 21.4% empty membrane. (F) The frequencies of the membrane coverage of all imaged HeLamPs are summarized. The number above each bar represents particle numbers for each bin.

Furthermore, STORM imaging unveiled a striking heterogeneity in the amount of membrane proteins carried by individual CNPs for both DCmPs and HeLamPs. While previous studies have determined that CNPs contain particles that are completely and partially coated with membrane proteins via indirect assessment,[21–23] STORM imaging provided an opportunity to assess the degree of membrane coating at the individual CNP particle level for the first time. We therefore quantified the membrane coverage of individual CNPs, defined as the percentage of the PLGA NP surface area covered by membrane proteins, and compared the coverage distributions of DCmPs and HeLamPs. (Fig 3A, 3B, 3D and 3F) For each particle cluster, we used the coordinates of the RhoB (PLGA) signals to reconstruct a two-dimensional core surface area, then estimated the fraction of that area colocalizing with WGA (membrane protein) signals by Monte Carlo Coverage Estimation.[33] Among all 4,697 imaged DCmPs, membrane coverage averaged about 22.7%, and about 29.0% of particles had coverage below 11%. The fraction of DCmPs decreased progressively as membrane coverage increased, and no DCmPs exceeded 81% coverage. (Fig 3D) In contrast, the membrane coverage of HeLamPs followed a normal distribution, with the majority of the 3,029 analyzed particles falling between 0.1% and 80%. The most frequent bin was 21-30%, accounting for about 21.1% of particles. No HeLamPs fell into the 91–100% coverage bin, and average coverage across HeLamPs was 37.8%. (Fig 3F) The differences in particle composition and membrane coverage distribution between DCmPs and HeLamPs are consistent with our finding that DCmPs have lower coating efficiency than HeLamPs. We attribute the observed coating performance differences to intrinsic membrane properties of the source cells, such as lipid composition or membrane fluidity, although the precise mechanism remains to be determined in future studies.[34] Compared with previously reported approaches, STORM imaging resolves membrane coating in substantially greater detail, enabling more rigorous structural characterization of CNPs and direct investigation of their structure–biofunction relationships. One caveat is that particles are captured on poly-L-lysine-coated slides, which may skew the observed CNP composition if membrane proteins bind poly-L-lysine more avidly than bare PLGA NPs. Future studies should also correlate STORM-derived membrane coverage with coating completeness measured by established indirect assays.[21–23]

Certain CNP biofunctions are driven by specific membrane proteins. For example, MHC proteins carried by DCmPs can engage and activate cognate T cells.[17,18] It is thus important to determine the composition and the quantity of function-relevant proteins for the assessment of CNP biofunctions. Considering the significant heterogeneity that we observed in membrane coating, we postulate that membrane protein composition of CNPs also varies greatly across CNP particles, which can not be assessed using conventional ensemble characterization approaches. We therefore adapted STORM imaging to characterize DCmPs with fluorescently labeled membrane proteins, focusing on MHC-I and the accessory proteins CD86 and ICAM-1, which are critical for T-cell engagement and activation.[35] Proteins were labeled with fluorescent antibodies, and individual particles identified algorithmically by fluorescence clustering. (Fig 4A) Across 5,166 DCmPs, 23.8% carried all three proteins, 34.5% MHC-I and ICAM-1, and 32.5% MHC-I and CD86. Notably, only 9.2% of particles were positive for ICAM-1 and CD86 alone, indicating that most particles carry MHC-I and potentially retain capacity for cognate T-cell binding. (Fig 4B) We then quantified relative protein abundance per particle using fluorophore localization counts. Although localization counts do not indicate exact protein copy number due to antibody labeling stoichiometry, they provide a reliable relative measure, enabling comparisons of protein levels across particles within a sample. (Fig 4C) Over 60% of DCmPs had fewer than 50 localizations for each protein. All three distributions were similarly right-skewed, with progressively fewer particles at higher counts. One important caveat is that DCmPs were identified solely based on membrane due to limited fluorescent channels (three). Because most membrane-positive particles are DCmPs, (Fig 3C) we think these abundance estimates remain relevant for DCmP coating characterization.

**Figure 4:**
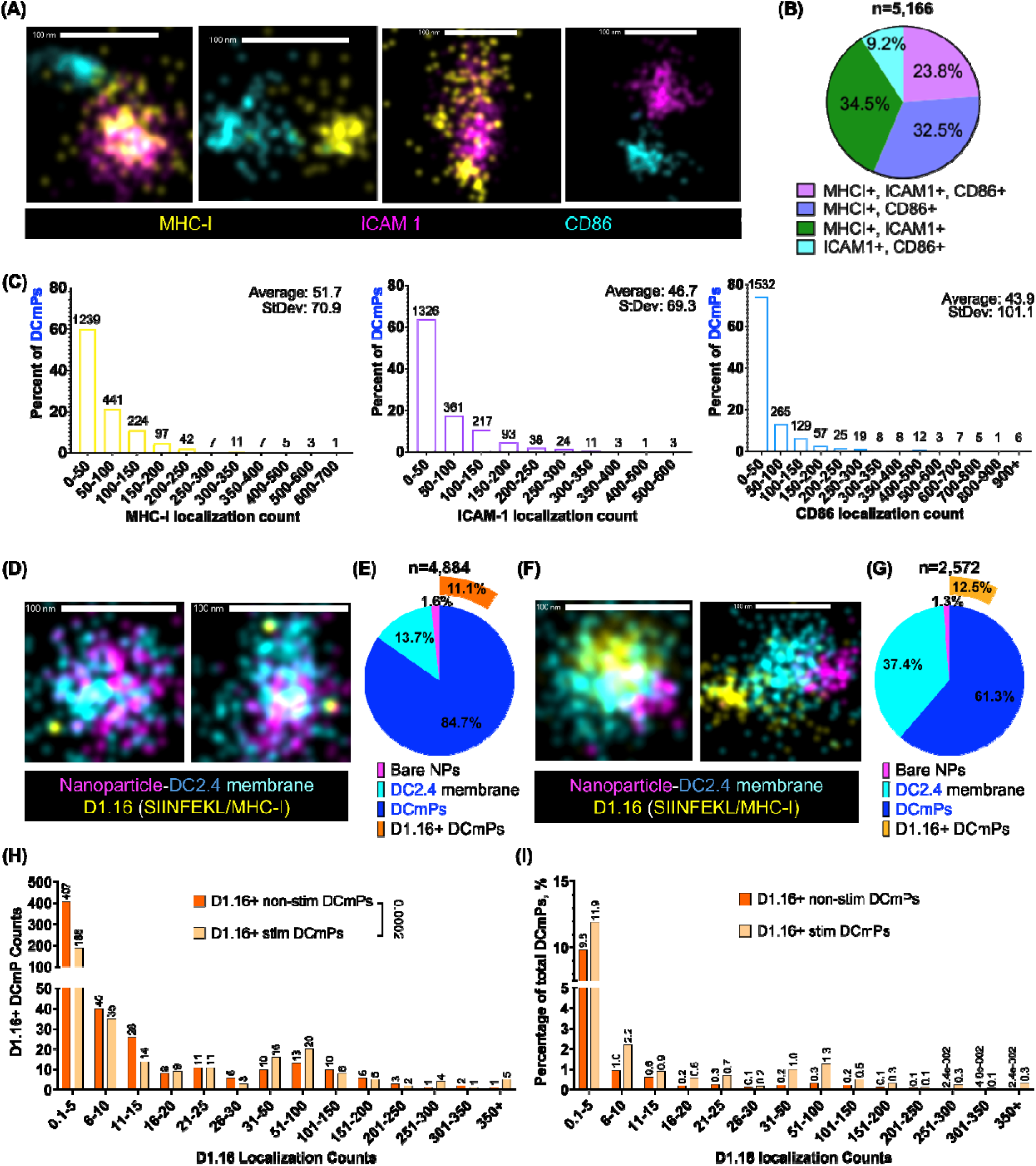
Quantification of membrane proteins carried by DCmPs using super-resolution imaging. (A) Representative super-resolution images showing the distribution of membrane proteins on individual DCmP particles. ICAM-1, MHC-I, and CD86 were labeled with fluorescent antibodies and imaged by STORM super-resolution microscopy. (B) Summary of the membrane protein composition on DCmPs. Of all 5,166 imaged DCmPs, 23.8% carried all three proteins, 32.5% were double-positive for MHC-I and CD86, 34.5% for MHC-I and ICAM-1, and 9.2% for CD86 and ICAM-1. (C) Distributions of localization counts for each protein across all individual particles. The average number of localization counts is 51.7 for MHC-I (StDev: 70.9), 46.7 for ICAM-1 (StDev: 69.3), and 43.9 for CD86 (StDev: 101.1). The number above each bar represents the corresponding particle numbers for each localization count range. (D) Representative images of non-stim DCmPs. (E) The particle composition of non-stim DCmPs as determined by STORM imaging. (F) Representative images of stim DCmPs. (G) The particle composition of stim DCmPs as determined by STORM imaging. (H) Distribution of D1.16 antibody localization counts for both non-stim DCmPs and stim DCmPs. The number above each bar represents total particle numbers for each localization count range. The statistical difference between the distributions of the two particle populations is determined using Chi-square test. (I) Distribution of D1.16+ particles among non-stim DCmPs or stim DCmPs. The number above each bar represents the frequency for each localization count range among membrane-coated particles.

We subsequently employed this STORM imaging approach to compare the abundance of antigen-specific MHC-I between DCmPs from OVA-stimulated DC2.4 cells and non-stimulated DC2.4 cells. After confirming SIINFEKL/MHC-I expression on stimulated DC2.4 cells (Fig S9), we stained both stim and non-stim DCmPs with D1.16 antibodies before STORM imaging. (Fig 4D and 4F) Of 4,884 non-stim particles, 84.8% were DCmPs, 1.6% bare NPs, and 13.7% free membranes; 11.1% were D1.16+ DCmPs. (Fig 4E) Among 2,572 stim particles, 61.4% were coated, 1.3% bare NPs, and 37.4% free membrane vesicles; 12.5% were D1.16+ DCmPs, slightly above the non-stim group. (Fig 4H) By flow cytometry, about 8% of non-stim and 18% of stim DCmPs were D1.16 positive. (Fig S10) These values is similar to the STORM results, confirming its utility for measuring protein abundance on CNPs. We then compared D1.16 localization-count distributions between stim and non-stim DCmPs. Both distribution were right-skewed toward low counts, but differed statistically (Chi-square, p=0.0002). (Fig 4H) Stim DCmPs showed higher frequencies across all count bins, confirming they carry more SIINFEKL/MHC-I complexes, which agrees with flow cytometry. (Fig 4I, S9, and S10) Therefore, STORM imaging reveals striking protein-composition heterogeneity among DCmPs and quantifies protein abundance on CNPs.

Because population composition here differed from our earlier measurements (Fig 3C), we asked whether batch-to-batch variation affected coverage for all stim and non-stim DCmPs (Fig S11). Average coverage was 13.5% (non-stim) and 25.4% (stim), the latter close to the 22.4% measured previously (Fig 3D). Both populations showed the right-skewed distribution seen in Fig 3D. Thus, despite compositional differences between batches, coverage varied little, suggesting membrane coverage is a more reproducible quality-control parameter for CNP manufacture. It is also worth noting that D1.16 antibodies bind non-stim DCmPs non-specifically, as seen in non-stim DC2.4 cells (Fig S9). Despite this caveat, the stim/non-stim comparison nonetheless remains informative for SIINFEKL/MHC-I levels.

Finally, we used a coculture assay to confirm that DCmPs preferentially bind cognate antigen-specific T cells. We have previously shown that DC2.4 cells predominantly express MHC-I, and that DCmPs from OVA-stimulated DC2.4 cells bind more B3Z cells (a CD8+ hybridoma with a TCR specific for SIINFEKL/MHC-I) than DOBW cells (a CD4+ hybridoma specific for OVA_323–339_/MHC-II) incubated separately.[18,36–38] To control for differences in culture conditions, we incubated a B3Z-DOBW coculture with fluorescently labeled DCmPs for 4 h and measured uptake by each cell type. (Fig 5A) The two populations were distinguished by CD4 expression (B3Z, CD8+CD4-; DOBW, CD8-CD4+). (Fig 5B and S12) After 4 h incubation, Over 18% of B3Z cells bound stim DCmPs (DiD+CFSE+), relative to only 2% of DOBW cells.. (Fig 5C and S13) B3Z binding to non-stim DCmPs was also significantly lower, indicating that cognate MHC/TCR interaction drives DCmP-T cell binding. Interestingly, a measurable fraction of both cell types also bound to bare NPs (DiD+CFSE-) during incubation with DCmPs, even though incubating cells with bare NPs alone produced no significant binding. This is consistent with our previous findings.[18] It may partially reflect a sensitivity limit of flow cytometry, in which particles bearing little membrane protein are still counted as bare NPs, which we are investigating further. Collectively, stim DCmPs bind T cells in an antigen-specific fashion even in the presence of non-cognate T cells.

**Figure 5:**
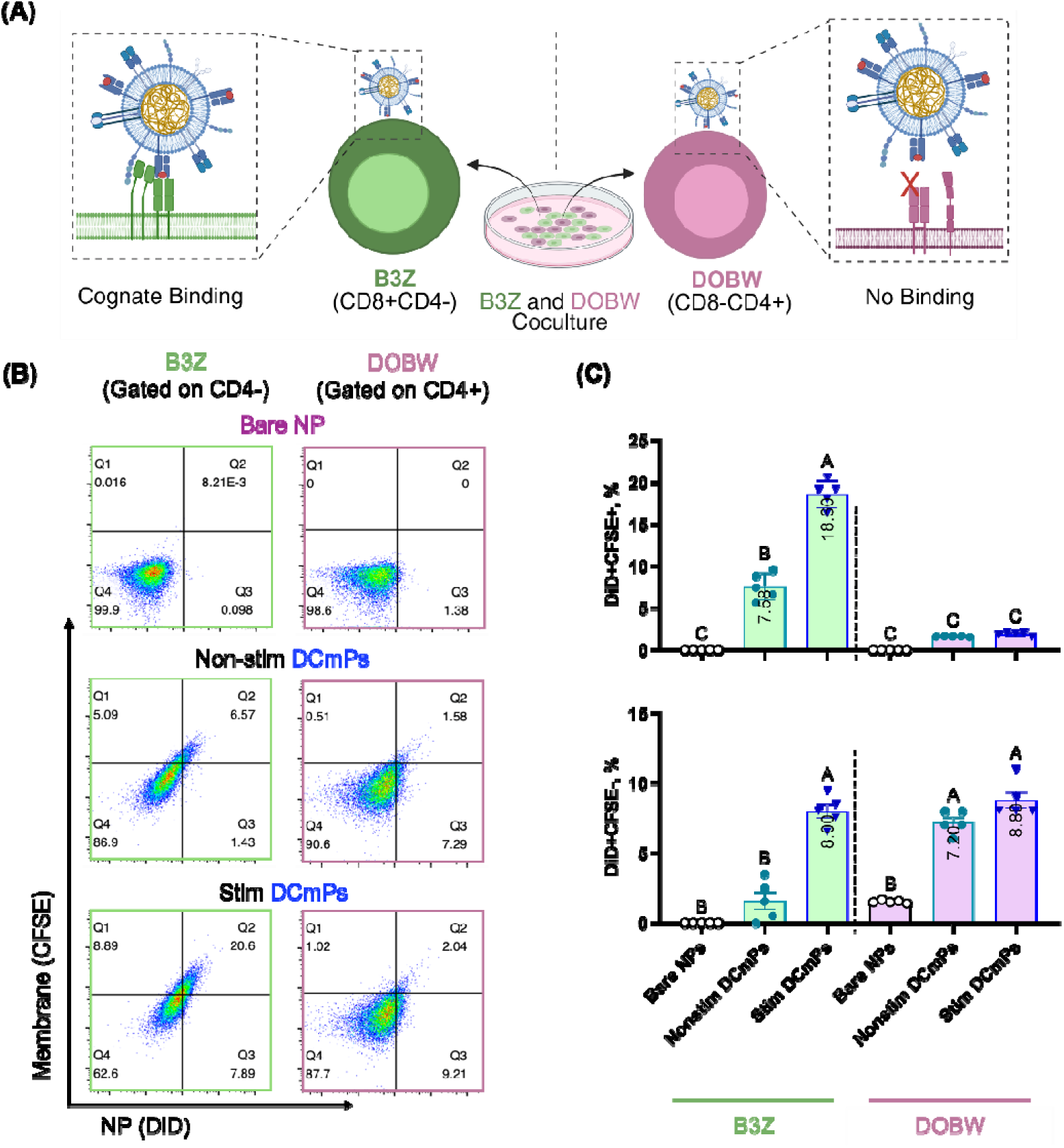
DCmPs preferentially bind to SIINFEKL-specific T cells in coculture. (A) Schematic of experimental design. SIINFEKL/MHC-I-carrying DCmPs preferentially bind to SIINFEKL/MHC-I-specific B3Z cells over OVA_323-339_/MHC-II-specific DOBW cells when incubated with the coculture of B3Z cells and DOBW cells. (B) Representative flow scatter plots of particle binding to cells. B3Z and DOBW cells are differentiated based on their expression of CD4. DCmPs are labelled with DiD (PLGA NP core) and CFSE (membrane proteins). (C) The frequency of fluorescent positive cells following incubation with bare NPs, non-stim DCmPs, or stim DCmPs, was assessed using flow cytometry. (Top: DiD+CFSE+; Bottom: DiD+) Statistical significance is determined using one-way ANOVA following Tukey’s multiple comparison test. Columns that do not share letters are statistically different from each other. N=5.

STORM super-resolution imaging has transformed structural biology, resolving organelle architecture and protein clustering at molecular level, and has recently been applied to the characterization of nanomaterials such as extracellular vesicles and lipid nanoparticles.[25–32] Here, we expand its capability to CNPs, achieving single-particle resolution unattainable by ensemble methods such as DLS and western blot. STORM confirmed that CNP populations comprise uncoated NPs, free membrane proteins, and coated particles, consistent with prior reports.[18,21–23] Critically, this approach further resolved the heterogeneity within the coated subpopulation itself, which is largely invisible to other conventional methods. For the first time, we quantified projected membrane coverage over individual NP surfaces and built full coverage distributions for both DCmPs and HeLamPs, which confirmed that DC2.4- and HeLa-derived membrane proteins behave distinctly during coating. Together, These findings establish single-particle membrane coverage as a quality metric for CNP characterization and a lever for optimizing and standardizing coating. Fluorescence localization counts also quantify protein abundance per particle, revealing in DCmPs marked particle-to-particle variation in surface protein content, establishing STORM as a platform for both spatial and molecular characterization of CNP coatings. A key limitation is that coverage analysis rests on 2D projections: each image collapses fluorescence from a particle’s proximal and distal surfaces into one focal plane. Z-axis resolution remains challenging given our particle sizes and current imaging platform. Reported coverage is therefore not absolute, but remains internally consistent and comparable across populations. Future work will quantify this projection error, potentially by correlative electron microscopy. Finally, a coculture assay confirmed that the coated membrane proteins remain functional: DCmPs engaged antigen-specific T cells in an MHC-dependent manner, consistent with the per-particle protein quantification by STORM and flow cytometry. More broadly, direct particle-level analysis via single-particle STORM imaging provides a rigorous quality metric, and facilitates CNP manufacture standardization and rational therapeutic design.

## Materials and Methods

### Cell Culture

All cell lines were cultured in a humidified incubator at 37°C with 5% CO_2_. Cells were passaged (by cell scraping) or cryopreserved upon reaching 80%+ confluency. DC2.4 cells (Merck, ref# SCC142) are cultured in RPMI 1640 medium (Thermo Fisher Scientific, ref# 21870-076), supplemented with 10% Fetal Bovine Serum (Thermo Fisher Scientific, ref# A5670801), 1% 1X Pen Strep (Thermo Fisher Scientific, ref# A5873601), 1% 1X L-glutamine (Gibco, ref# 25030-081), 1% 1X MEM NEAA (Gibco, ref# 11140-050), 2.5% 25mM HEPES (Corning, ref# 25-060-CI), and 0.0054% β-Mercaptoethanol (Gibco, ref# 21985-023). HeLa cells were kindly gifted from Dr. Xiaoran Hu (Department of Chemistry, Syracuse University, Syracuse, New York 13210, United States). HeLa cell medium is formulated with DMEM (Gibco, ref# 11965-092), 10% Fetal Bovine Serum (Thermo Fisher Scientific, ref# A5670801), and 1% 1X Pen Strep (Thermo Fisher Scientific, ref# A5873601).

### DC2.4 Stimulation

DC2.4 cells stimulated with 0.3mg/mL of ovalbumin (Sigma-Aldrich, ref# A5503-1G) and 50ng/mL of lipopolysaccharide (Novus Biologics, ref# NBP22529510) in either T75 or T175 flasks. Stimulation occurred for 22-26 hours. Cells then washed with PBS, centrifuged at 500 x g for 5 minutes at RT, and collected. Cells stored at -80°C in 1X TRIS-buffered saline (TBS), pH 7.4 (Thermo Fisher Scientific, ref# AAJ60764K2) with protease inhibitor cocktail (Thermo Fisher Scientific, ref# 78442) before membrane isolation.

### Membrane Isolation

Cells were thawed in 37°C water bath and transferred to an 8mL Dounce homogenizer on ice. Cells were mechanically disrupted by 40 passages using the loose pestle followed by 40 passages using the tight pestle. The lysate was centrifuged at 3000 G for 10 minutes at 4°C. Supernatant was collected into an ultracentrifuge-safe 15mL tube (Beckman Coulter, ref#NC9194790). Cell pellet was resuspended in 1xPBS with protease inhibitor and treated with a second round of 40 loose pestle and 40 tight pestle mechanical disruption passages. The resulting lysate was centrifuged and pooled with the previously collected lysate in the 15mL ultracentrifuge tube. Pooled lysates were centrifuged in an ultracentrifuge (Thermo Scientific wX+ Ultra Series, TH-641 rotor) at 100,000 G for 60 minutes at 4°C. The resulting pellets were resuspended in 1X PBS with 0.05% Tween-20 (PBST). Sample protein concentration was measured by Bradford (595 nm wavelength) and stored at -80°C before coating.

If CFSE stained membrane is used, cells were stained with CellTrace CFSE stain (Invitrogen, ref# C34554) following manufacturer recommended protocol before membrane isolation.

If Wheat Germ Agglutinin (WGA) labelled membrane proteins are used, cells were treated with Alexa Fluor 488 conjugated Wheat Germ Agglutinin (Invitrogen, ref# W11261) at 5.0ug/mL and incubated times at 37°C for 10 minutes. Labelled cells were then washed with PBS before membrane isolation.

### Fabrication of PLGA Nanoparticles

Poly(lactide-co-glycolide) (PLGA, 50:50, IV 0.6 dL/gl) (Polysciences, Cat# 23986) was dissolved in acetonitrile at 1mg/mL concentration. In a typical synthesis, 1 mL of PLGA solution was added dropwise into 13 mL of 0.7% polyvinyl alcohol (PVA) under constant 600 rpm stirring for nanoprecipitation for 4 hours and left overnight at 200 rpm. Nanoparticles were resuspended in deionized water and filtered through a 100kDa Centricon filter tube (Thermo Fisher Scientific, ref# UFC910024) at 4000 G at room temperature. After 5-time DI-water wash, PLGA NPs were collected and freeze-dried. For RhoB (Thermo Scientific ref# A1357218) labeling, 0.005% w/v RhoB was added to the PVA aqueous phase. If DiD (Invitrogen, ref# D307) dye is used, DiD DMSO solution was added to the PLGA/ACN organic phase solution at 0.0375 (w/w)% prior to nanoprecipitaion. Precautions were taken to minimize light exposure when dyes were used.

### Fabrication of Membrane Coated PLGA NPs

A combination process of sonication and extrusion was performed to fabricate the coated PLGA NPs. 500 µg of NPs and isolated membrane proteins were combined at a 1:1 weight ratio (µg dry weight NP: µg protein) in 1mL of PBS and sonicated for 5 minutes in sonication bath, followed by centrifugation at 10,000 G for 5 minutes at 4° C to remove unbound proteins. The pellets were resuspended in 1mL of PBS and extruded through a 200 nm polycarbonate membrane (Avanti, ref# 6100061EA) for 11 times using an Avanti Polar Lipid Extruder (Avanti, ref# 6100001EA). Protein coating quantity was determined via Bradford assay (Thermo Fisher Scientific, ref# 23238).

### Cellular Uptake

2 × 10 DC2.4 or HeLa cells were seeded onto 96-well plates and incubated with each particle formulation at 100 μg/mL (based on PLGA weight) for 4 hrs at 37 °C with 5% CO (n = 5). Cells were then washed three times with cold flow buffer (PBS plus 2% FBS) and then stained with FVD, before fixing with 4% paraformaldehyde for flow cytometry analysis using a BD LSR Fortessa analyzer.

For confocal microscopy, 2 × 10 cells were seeded in sterile 4-well chamber slides and incubated with particles under the same conditions. Images were acquired using a Leica DMi8 CLSM. The colocalization analysis of RhoB-labeled PLGA and CFSE-labeled membrane was conducted using Fiji (ImageJ) following Manders’ coefficients. Four independent images were analyzed.

### ZetaView

PMX130 Mono instrument and software version: ZetaView (version 8.06.01 SP1) was used to obtain size and zeta measurements. Particle samples were prepared at 1mg/mL (based on PLGA weight) and further diluted by 4000-fold in PBS. ZetaView cell primed and washed after quality checks, and between each sample analysis. Sample volume added to ensure appropriate cell coverage. 11 cell sections analyzed per sample. ZetaView used at 25°C. Samples were analyzed using protocol EV520.

### Dynamic Light Scattering (DLS)

Zetasizer instrument and ZS Xplorer software were used to obtain measurements. Samples were prepared at 1mg/mL in PBS and added to a cuvette (DTS0012 or DTS1070). DTS0012 used exclusively for size measurements. DTS1070 used for zeta potential and size reads. Three technical reads per independent samples were performed.

### SDS-PAGE and Western Blotting

Samples are prepared at a total volume of 50µL, a mixture that included 2x sample buffer (Novex, ref# LC2676), including particle samples, proteins and deionized water as needed. Protein samples denatured on a heat block at 75°C for 5 minutes, before being loaded into 4-12% Tris-Glycine Gel (Invitrogen ref# XP04120BOX) with 1x running buffer. 5-8µL of standard protein ladder (Thermo Fisher Scientific, ref# 26619) was also loaded along with protein samples. Gel electrophoresis was run at 80V for 20 minutes initially, then at 100 V until the gel was resolved (approximately 60-80 minutes in total). For SDS-PAGE analysis, gels were stained with Blue Stain (Thermo Fisher Scientific, ref# 24590) for 1 hour, shaking at 70rpm. The gel was then destained by three deionized water washes.

For western blotting, the gel and a 0.45 µm membrane (Cytiva, ref# 10600002) were placed in a transfer tank system for protein transfer at 75V for 75 minutes on ice. The membrane was then blocked with 5% milk in TBS with 0.05% TWEEN20 (TBST) for 1 hour at room temperature, before being incubated with primary antibodies for 24 h at 4° C on a shaker. Primary antibodies include anti-MHC Class I (Cell Signaling Technologies, Cat# 76828), anti-CD86 (Cell Signaling Technologies, Cat# 19589), anti-ICAM-1 (Thermo Fisher Scientific, Cat# PI701254), and Na+/K+ ATPase (Cell Signaling Technologies, Cat# 3001S). Secondary HRP-conjugated goat anti-rabbit antibodies (Invitrogen, ref# 32460) were then added for imaging. Images were obtained by a BioRad ChemiDoc MP Imaging System (software version 2.4.0.03), and analyzed using BioRad ImageLab software.

### STORM Super Resolution Microscopy

Chamber slides (Part# 800-00095) were coated with Poly-L-Lysine (EMD Milllipore cat#A-005-C) for 15 minutes to facilitate sample attachment, with excess Poly-L-Lysine removed with PBS wash. 10 µL of sample (50 µg of protein) was added to the slide and incubated for 50 minutes, followed by PBS wash twice. The slide was then incubated for 45 minutes with the following fluorescent antibodies in the presence of Fc block (BD Biosciences, Cat# 553142) as needed: anti-MHC Class I AF488 (Cell Signaling Technologies, Cat# 567696), anti-MHC Class I DyLight550 (Novus, Cat# NB10065938S), anti-ICAM-1 AF647 (Invitrogen, Cat# A15397), anti-CD86 FITC (BD Biosciences, Cat#BDB553691), or AF647 Anti-Mouse H-2K[b]/SIINFEKL (BD Biosciences, Cat# 570092). All antibodies were used at a 1:50 dilution. After PBS wash twice, slides were imaged on ONI Nanoimager microscope (NimS-Mark III) with AutoEV program (CSA 1.3.7). Images were processed and clusters algorithmically determined by the “EV Profiling” protocol provided in ONI CODI software (CSA 1.3.7).

For EV Profiling analysis, Drift correction was performed, followed by noise filtering via restricting the fluorescent frame index range and background removal using a blank control. During DBSCAN analysis, the minimum density of the clusters was set at 15, and minimum number of localizations per cluster was set at 30. RhoB and WGA channels were not merged. MHC-I, ICAM1, and CD86 channels were merged. Cluster filtering was then performed on DBSCAN results. Cluster area (nm^2^) and radius of gyration (nm) were set based on coated NP measured size by ZetaView. Circularity range set to 0.8-1.0. Localization density was set to 0.001-1.0nm^2^. All other parameters stayed standard for ONIs EV Profiling software.

Membrane coverage of PLGA nanoparticles was quantified using a custom Python program. Clustering output from Alto.Codi.Bio (cluster length in nm and x, y coordinates) was used to construct a 2D cluster area for each particle. Drift-corrected and filtered single-molecule fluorescence points were then mapped onto membrane positions within each cluster boundary, and surface coverage was estimated by Monte Carlo simulation (100,000 independent events per cluster). Colocalization between the membrane and NP core signal was scored as present or absent at each surface point where a Monte Carlo event was analyzed. The fraction of events scored as non-colocalized was taken as the estimate of uncoated surface area. Bare nanoparticles and free membranes were excluded from the analysis.

Protein localization distributions were calculated using the EV Profiling method. Distributions were determined individually for each channel based on positive signal, and percentages were calculated relative to the entire imaged population.

### Statistical Software and Figure Generation

Data processing and figure generation were performed in Python (Matplotlib) using the Spyder IDE (v6.0.7; Python 3.11.12) and in GraphPad Prism 11. Flow Cytometry results processed in FlowJo 10.10.0. Figures 1A and 5A were created with BioRender. All statistical tests used a significance threshold of p < 0.05; the specific test for each figure is noted in its caption.

### ASSOCIATED CONTENT

## Supporting Information

The following files are available free of charge.

Membrane protein SDS-PAGE gel images, DLS measurements of CNPs, CNP stability analysis, flow cytometry gating strategies, additional confocal images of cellular uptake for both HeLa and DC2.4 cells, flow cytometry analysis of SIINFEKL/MHC-I level in DC2.4 cells and DCmPs, additional analysis of membrane coverage distribution for both non-stim DCmPs and stim DCmPs, DOBW and B3Z cell surface marker confirmation, and gating strategy for DCmP-T cell binding experiments.

### AUTHOR INFORMATION

## Present Addresses

†Present Address of Sao Puth: The Global Health Research Center, University of Health Sciences, Khan Daun Penh, Phnom Penh, 12000, Cambodia

## Author Contributions

J. B. W., A. Z., S.P. and Y. W. conceived the project, designed the experiments, analyzed data, and wrote the manuscript. J. B. W., A. Z., S. P., S. S. J., and R. C. performed particle synthesis and characterization. J. B. W. performed STORM imaging and analysis. S. P. and A. Z. validated DC stimulation, performed confocal imaging, and flow cytometry experiments. A. M. H. performed flow cytometry experiments. A. J., J. Z., J. L. H. provided resources. Y.W. provided resources and supervision. All authors reviewed and approved the manuscript.

## Funding Sources

This research is supported by the Syracuse University Startup Fund, Syracuse University G2G Fund, and NIH NIAID R21AI196694.

## Notes

The authors declare the following competing financial interest(s): Y. W. is listed as an inventor on pending patent applications. All other authors declare no conflicts of interest.

## ACKNOWLEDGMENT

We extend our sincere appreciation to Dr. James Moon at the University of Michigan for kindly providing B3Z cells and to Dr. Clifford Harding at Case Western Reserve University for the generous gift of DOBW cells. The illustrations were created in BioRender (https://BioRender.com/u08ltfd).

