## Supplemental Files for "Single-Particle STORM Imaging Quantifies Coating Heterogeneity Among Cell Membrane-Coated Nanoparticles"

AUTHOR ADDRESS

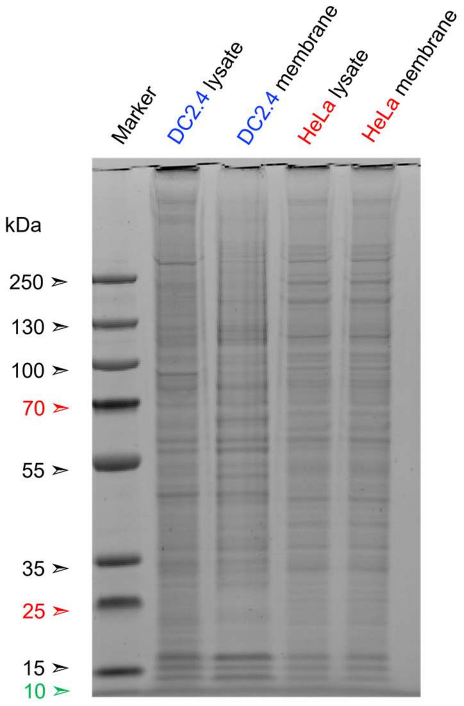


**Figure S1: Membrane protein characterization.** Protein profile comparison between source cell lysate and membrane proteins of source cells, according to SDS-PAGE.

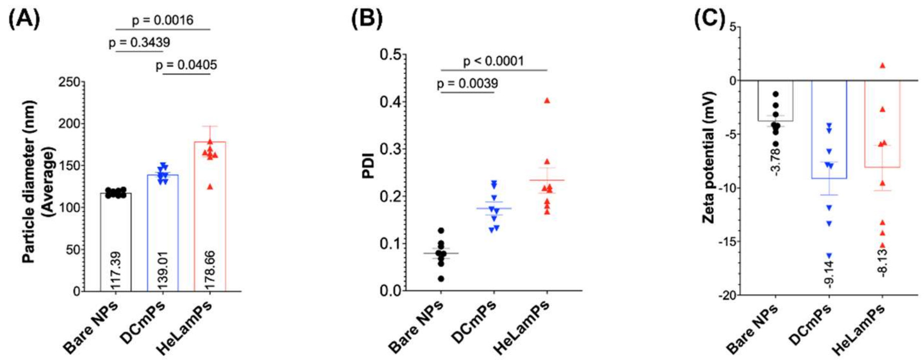


**Figure S2. CNP Characterization via DLS Particle diameter** (A) polydispersity (PDI), (B) zeta potential (C) of bare NPs, DCmPs, and HeLamPs are determined by dynamic light scattering (DLS). Statistical significance is determined via one-way ANOVA, n=8.

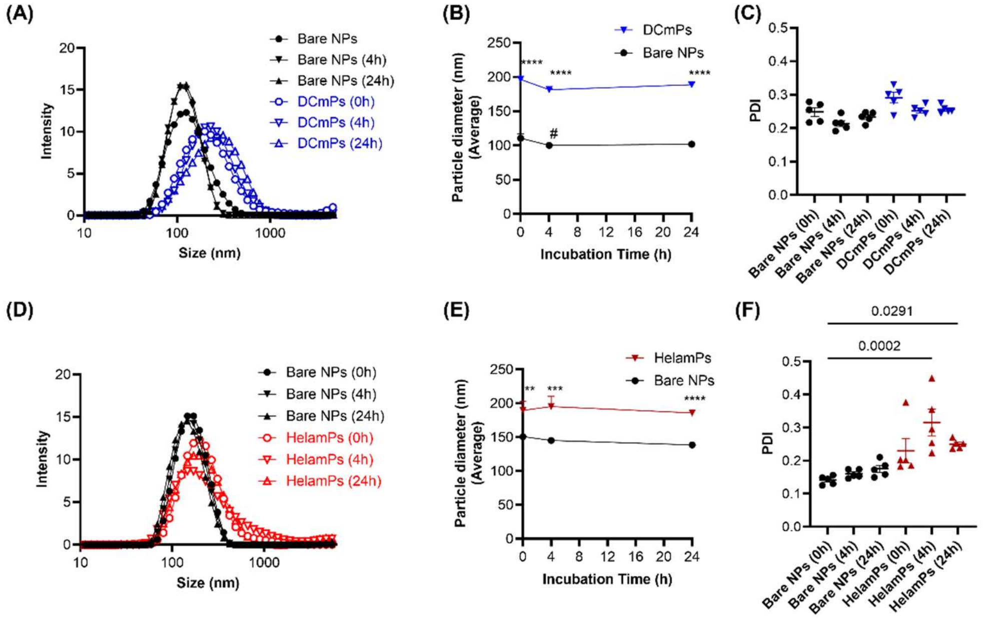


**Figure S3. Both DCmPs and HeLamPs are colloidally stable in PBS for 24 h**. (A) DCmPs and bare NPs are incubated in serum-free PBS buffer with their diameters and polydispersity index (PDI) measured via DLS at 0, 4, and 24 hours. The average size distributions of bare NPs and DCmPs at each time point are shown, N=5; (B) The changes in z-average diameters of bare NPs and DCmPs over 24 hours. Statistical significances are determined with two-way ANOVA with Tukey's multiple comparisons test. **** represents statistical significant difference between DCmPs and bare NPs at corresponding time points ($p<0.0001$); # represents statistical significant difference when measured diameters are compared to the 0 h time point for each particle. ($p<0.05$). N=5. (C) Polydispersity index (PDI) of bare NPs and DCmPs over the experimental period. N=5. (D) HeLamPs and bare NPs are incubated in serum-free PBS buffer with their diameters and PDI measured via DLS at 0, 4, and 24 hours. The average size distributions of bare NPs and DCmPs at each timepoint are shown, N=5; (E) The changes of z-average diameters of bare NPs and HeLamPs over 24 hours. Statistical significances are determined with two-way ANOVA with Tukey's multiple comparisons test. *

represents statistical significant difference between HeLamPs and bare NPs at corresponding time points (** p<0.01, *** p<0.001, ****, p<0.0001), N=5;  (F) PDI of bare NPs and HeLamPs over experimental period. Statistical differences are determined with one-way ANOVA with Tukey's multiple comparisons test. N=5.

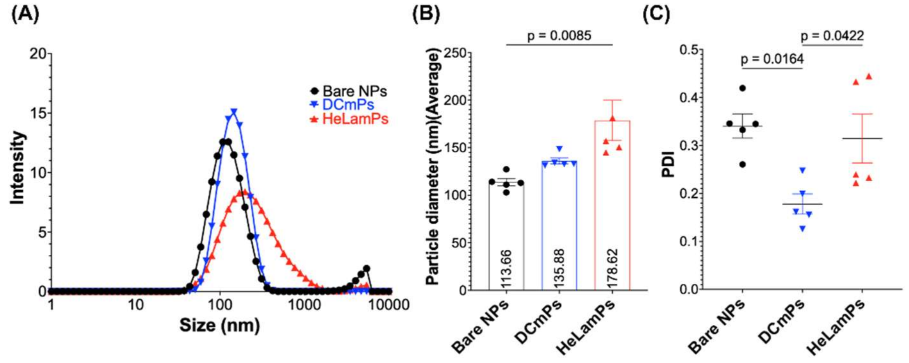


**Figure S4. Characterization of fluorophore-labelled membrane-coated nanoparticles.** Both DCmPs and HeLamPs are produced using RhoB-encapsulated PLGA NPs and CFSE-labelled membrane proteins. (A) Size distribution (B) Z-average diameters of bare NPs, DCmPs and HeLamPs (C) PDIs of bare NPs, DCmPs, and HeLamPs. Statistical significance is determined using one-way ANOVA with Tukey's multiple comparison test. N=5.

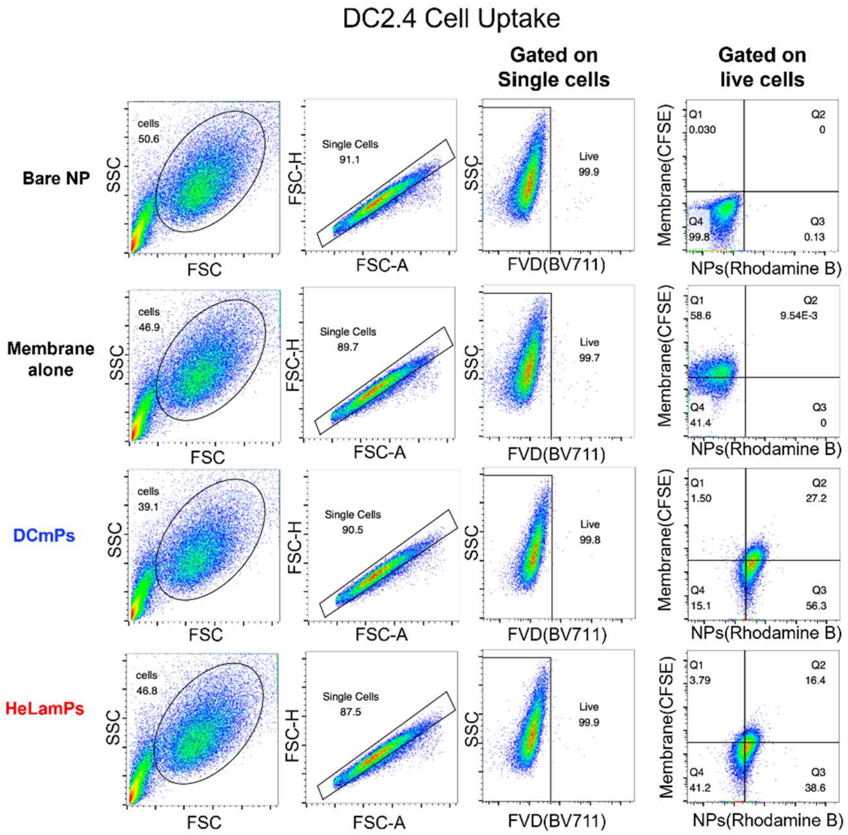


**Figure S5. Representative flow plot and gating strategy for DCmP and HeLamP uptake by DC2.4 cells at 2 hours.**

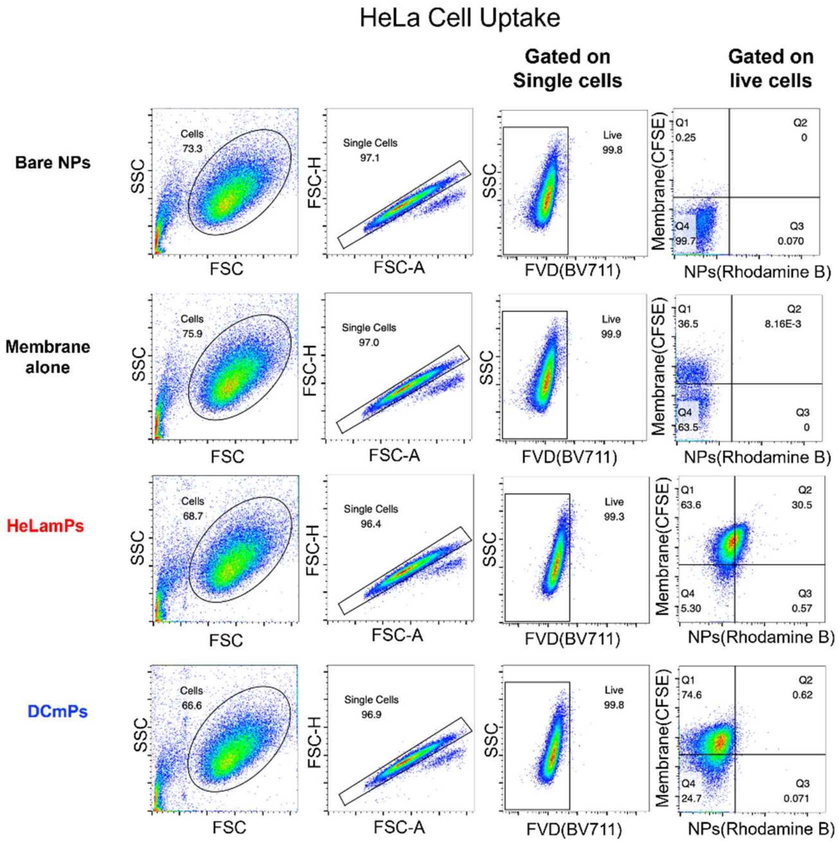


**Figure S6. Representative flow plot and gating strategy for DCmP and HeLamP uptake by HeLa cells at 2 hours.**

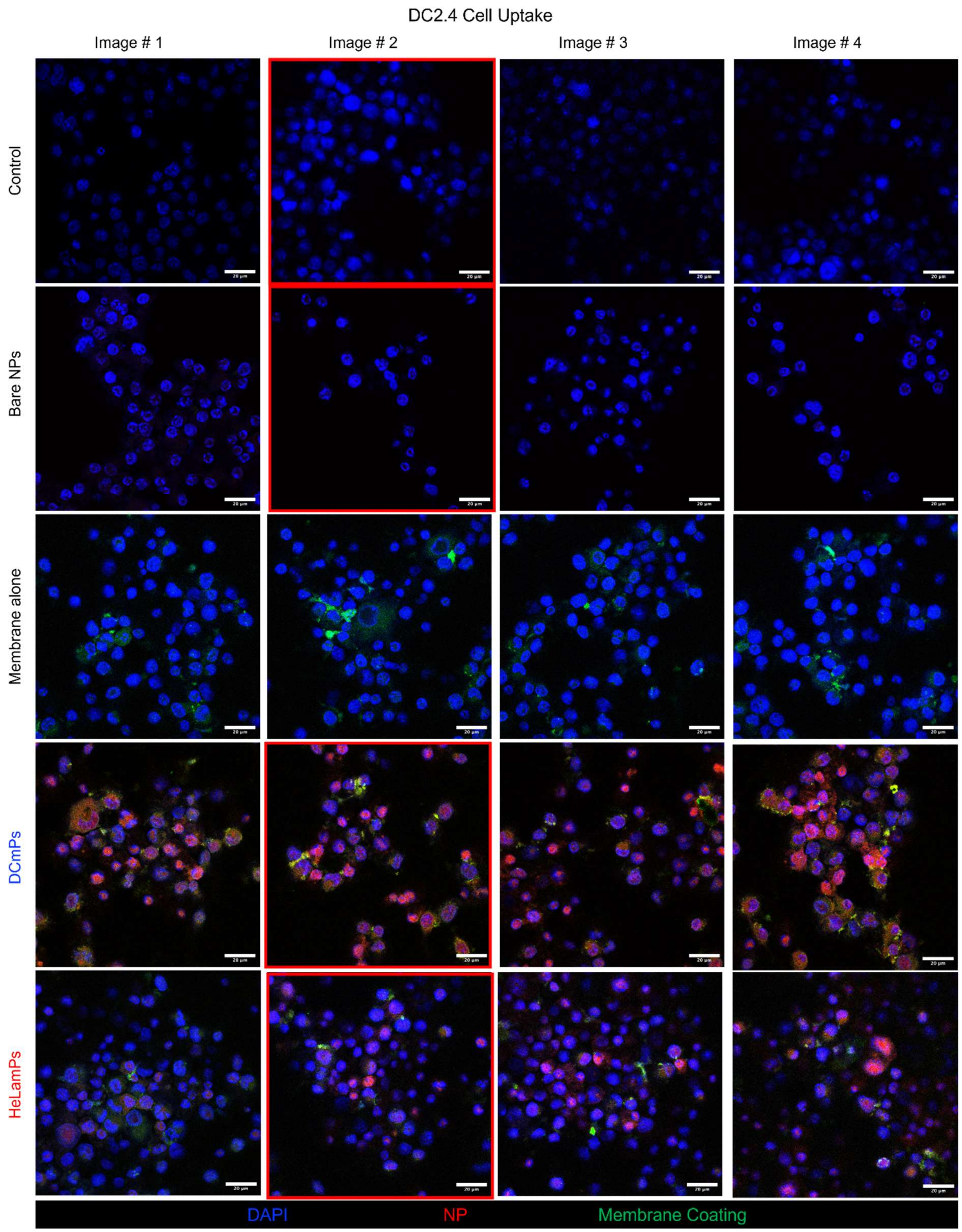


**Figure S7: Confocal images of control, bare NPs, membrane alone, DCmP, and HeLamP uptake by DC2.4 cells after 4 hour incubation.** Membrane coating is labelled with CFSE (green), RhoB-encapsulated NPs are shown in red. Nuclei are stained with DAPI.

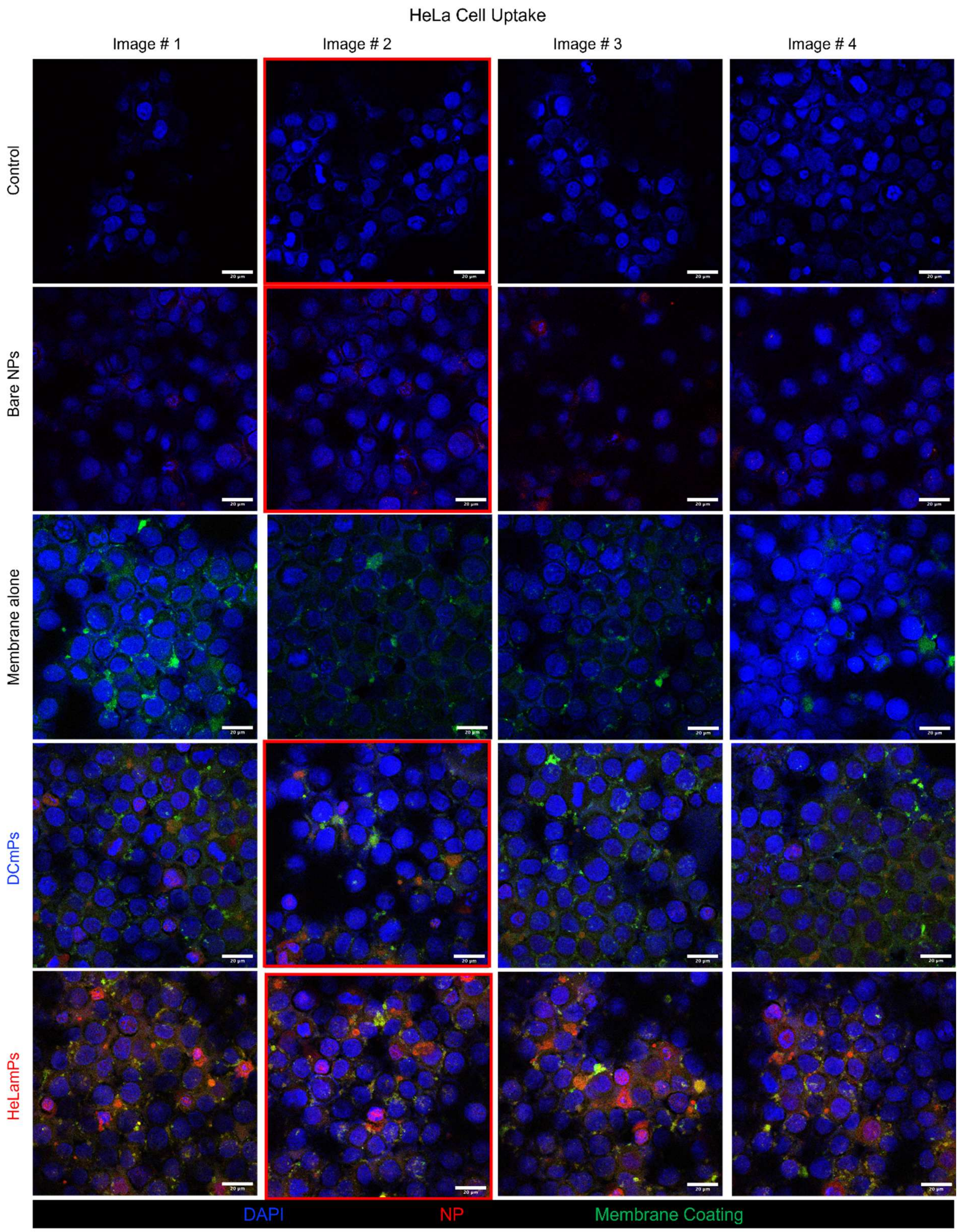


**Figure S8: Confocal images of control, bare NPs, membrane alone, DCmP, and HeLamP uptake by HeLa cells after 4 hour incubation.** Membrane coating is labelled with CFSE (green), RhoB-encapsulated NPs are shown in red. Nuclei are stained with DAPI.

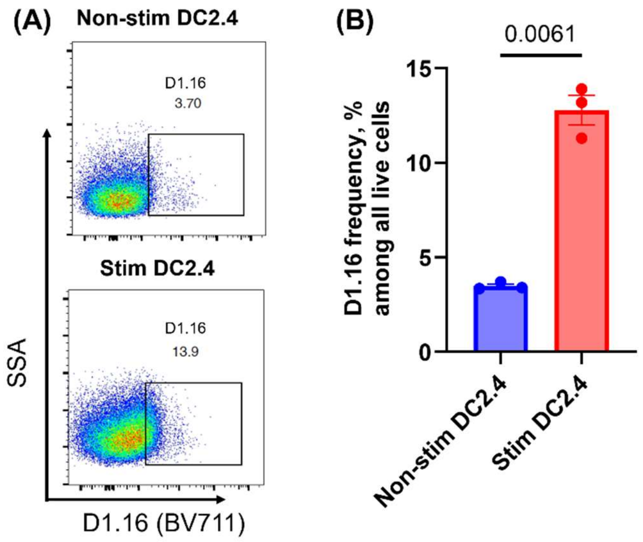


**Figure S9. Confirmation of upregulation of SIINFEKL/MHC-I in OVA-stimulated DC2.4 cells via flow cytometry**. (A) Representative flow histograms of D1.16 staining on DCmPs. (B) Summary of D1.16+ DC2.4 cell frequency with and without OVA stimulation. Statistical significances are determined using an unpaired t-test with Welch's test. N=3

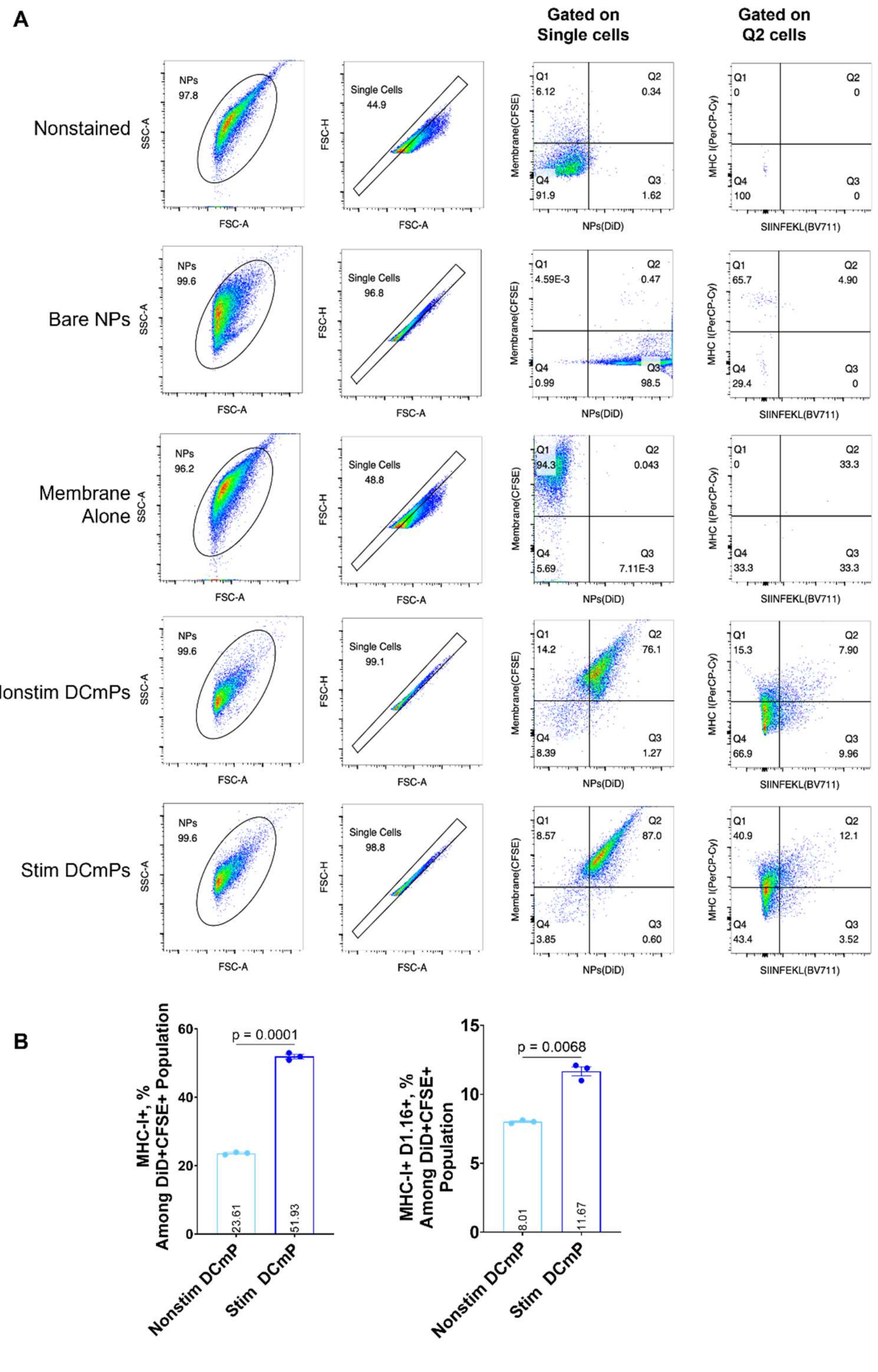


**Figure S10. SIINFEKL/MHC-I analysis on nonstimulated and stimulated DCmPs**. (A): Representative flow plots and gating strategy. (B): Frequencies of MHC-I+ and MHC-I+ and 25-D1.16+ populations among DCmPs coated with membrane proteins from nonstimulated and OVA-stimulated DC2.4. Statistical significance is determined using unpaired student t-test with Welch's test. N=3

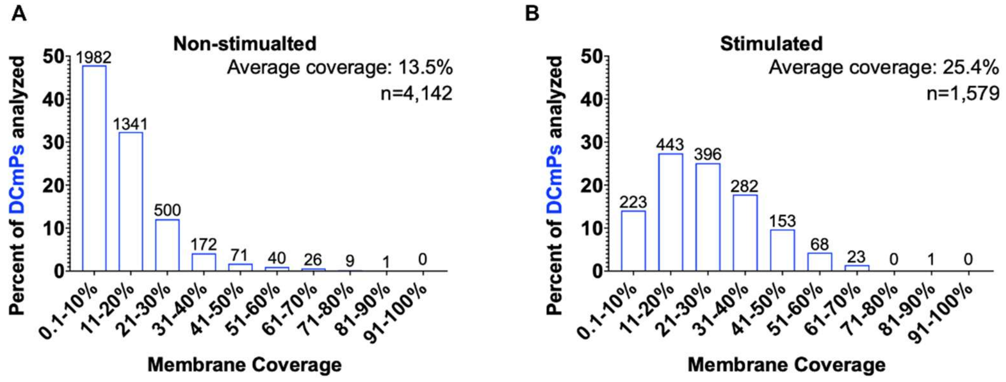


**Figure S11. Single-particle membrane coverage distribution.** The membrane coverage of non-stim DCmPs from Fig 4E **(A)** and stim DCmPs from Fig 4G **(B)** are determined using STORM imaging. The number above each bar represents particle numbers for each membrane coverage range.

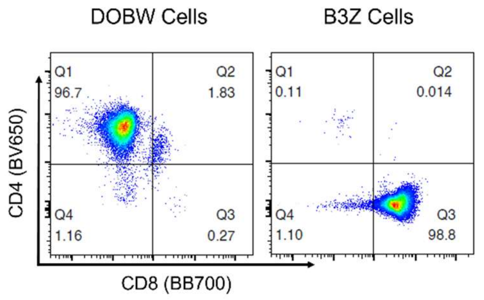


**Figure S12: DOBW and B3Z Surface Maker Characterization.** DOBW and B3Z cells show distinct surface markers. DOBW cells are CD4+CD8-, and B3Z cells are CD4-CD8+.

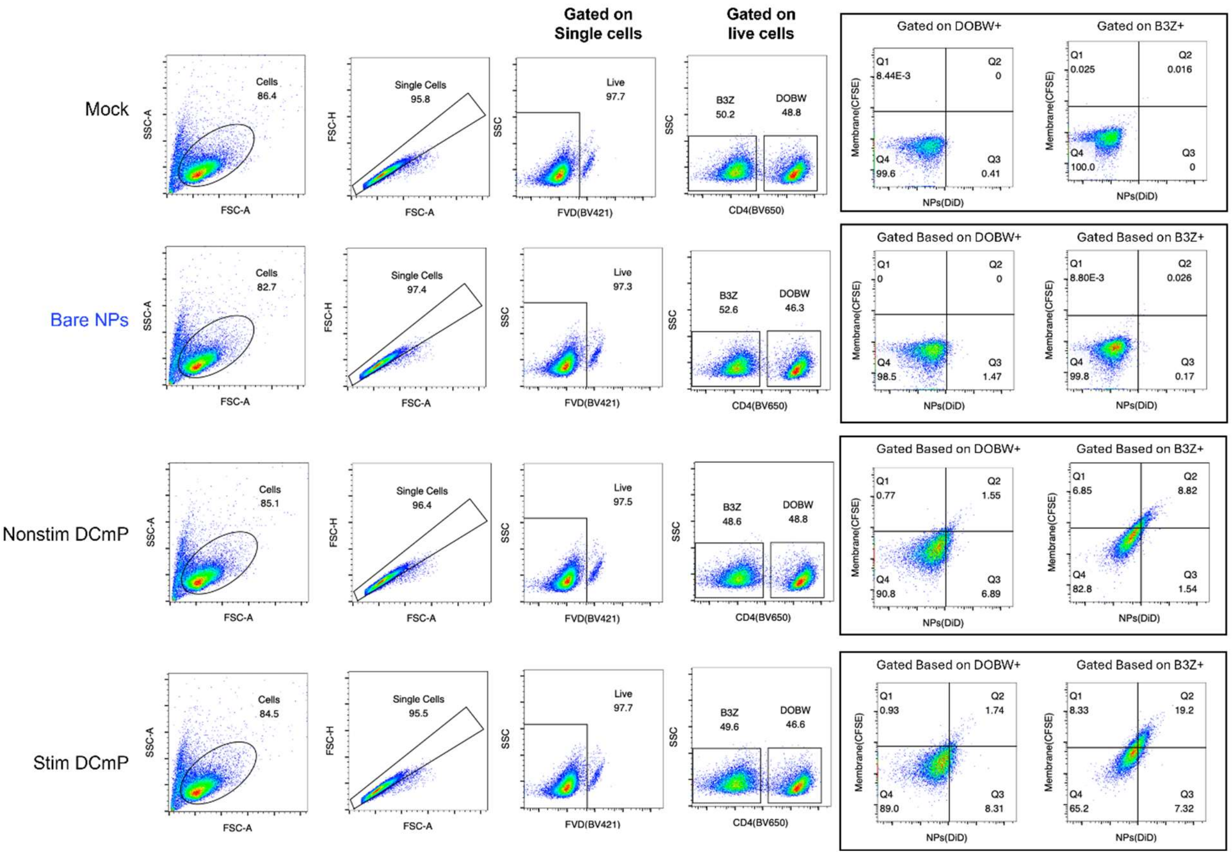


**Figure S13: The gating strategy and representative flow plots of cellular binding to DCmP after 4-hour incubation in B3Z and DOBW coculture.**
